# SLR-superscaffolder: a *de novo* scaffolding tool for synthetic long reads using a top-to-bottom scheme

**DOI:** 10.1101/762385

**Authors:** Lidong Guo, Mengyang Xu, Wenchao Wang, Shengqiang Gu, Xia Zhao, Fang Chen, Ou Wang, Xun Xu, Guangyi Fan, Li Deng, Xin Liu

## Abstract

Synthetic long reads (SLR) with long-range co-barcoding information have been recently developed and widely applied in genomics researches. We proposed a scaffolding model of the co-barcoding information and developed a scaffolding tool with adopting a top-to-bottom scheme to make full use of the complementary information in SLR datasets and a screening algorithm to reduce negative effects from misassembled contigs in an input assembly. In comparison with other available SLR scaffolding tools, our tool obtained the best quality improvement for different input assemblies, especially for those assembled by the next-generation sequencing reads, where the improvement of contiguity is about several hundred-folds.

## Background

Single tube long fragment reads (stLFR) library [1], as one of synthetic long reads (SLR)technologies [2-5], has been recently developed for effectively co-barcoding short next generation sequencing (NGS) reads from the same long DNA fragment (LDF). Since the co-barcoding information for each LDF is recoverable according to the same barcode shared by reads from the specific LDF, it can be applied to haplotyping [2,4-7], structural variation detection [8-10] and *de novo* genome assembly [11-17]. Like the previous whole genome shotgun strategy of sequencing for the BAC [18] or Fosmid library [19], a stLFR library can also retain the long-range information of LDFs, but be more cost-effective to be widely applied in genome researches.

As other SLR technologies [2-5] the co-barcoded reads coverage depth of a single LDF in a stLFR library is too low to implement a direct assembly through a two-stage process [20] where LDFs are firstly assembled by reads with the same barcode, and then the genome is further assembled by the pre-assembled LDFs. However, the enhancement in the LDF coverage depth of genome could overcome the weakness of current SLR technologies.

To utilize the co-barcoding information, there have been several genome assembly tools previously designed for each specific SLR technology. For contiguity preserving transposition sequencing (CPT-seq) reads, Adey, et al., developed fragScaff to conduct scaffolding using the minimum spanning tree (MST) algorithm on a scaffold graph constructed by the co-barcoding information[11]. Their results of the whole human genome show that the improvement of the input assembly with a large NG50 of about 100 kb is significantly better than that with a small NG50 of about 10kb. For SLR sequencing technology from Illumina [3], Kuleshov et al., developed Architect by heuristically removing spurious edges in a scaffold graph constructed by combining the co-barcoding and paired-end (PE) information [12]. In their tests of species with small genome sizes, the improvement of final assembly is also dependent on the contiguity of the input assembly. For 10X Genomics Chromium technology (10XG-linked reads) [5], Warren, et al., developed a scaffolder with two versions ARCS [14] and ARKS [15] based on LINKS [21], which is a scaffolder initially designed to use long reads sequenced by single molecular real time sequencing (SMRT) technologies. Their results show that ARKS can make significant improvements for the input assembly with a very large NG50 such as 4.6 and 14.7 Mb. In addition, Weisenfeld, et al., from 10X Genomics, developed a *de novo* assembler named Supernova for the 10XG-linked reads, which could generate diploid genomes from raw reads [13]. Recently, a universal assembler CloudSPAdes was developed by Tolstoganov, et al., for SLR datasets to combine an assembly graph with the co-barcoding information based on SPAdes assembler [17, 22]. Both Supernova and CloudSPAdes do not provide independent module for scaffolding, but the co-barcoding information was also used for scaffolding according to the description of their algorithms. The dependence of efficiency of using the co-barcoding information on the quality of an input assembly cannot be directly evaluated for these two tools. For standalone scaffolders, an input assembly with long contiguity is required to make highly efficient use of the co-barcoding information. Meanwhile, the SLR datasets with different characters were used in the developments of those tools. Thus, it is still a challenge to develop a robust scaffolder with a low requirement of the quality of an input assembly to improve different draft assemblies by using various SLR datasets.

Here we proposed a general scaffolding model for using the co-barcoding information of a SLR dataset and developed a standalone scaffolder (SLR-superscaffolder) with using stLFR reads. As a standalone scaffolder, only a draft assembly (contigs or scaffolds) and a SLR dataset are required as input. The final assembly is dependent on the quality of both the PE and co-barcoding information and also on the quality of an input assembly. In our tool, we adopted an overall top-to-bottom scheme to make full use of the complementary information in co-barcoded reads and to reduce the contiguity requirement of input assemblies and a screening algorithm to reduce negative effects of non-ideal seed contigs in an input assembly. In addition, our tool was designed with a high degree of modularity, where the order, orientation and gap size were determined one by one. All the above considerations may make our tool more robust to different input.

In our development of SLR-superscaffolder, properties of the PE and co-barcoding information in stLFR dataset were analyzed by aligning reads to the reference genome of the human whole genome for cell line NA12878 (HG001). The results demonstrate that the number of LDFs per barcode is significantly small and the distribution of insert size of PE reads is irregular. According to the scaffolding model, different types of correlation between two sequences were analyzed, and Jaccard Similarity (JS) was used to construct a co-barcoding scaffold graph in SLR-superscaffolder. To benchmark different scaffolders including SLR-superscaffolder, fragScaff, Architect and ARKS, the combinations of stLFR reads with other sequencing datasets, such as PCR-free NGS reads and ONT reads, were tested together for HG001. The benchmark results show that the scaffolds assembled by SLR-superscaffolder are with longer contiguity and higher accuracy than others for draft assemblies assembled by using NGS reads. Considering the generality of stLFR reads, SLR-superscaffolder would have great potential to be a universal scaffolder for various SLR datasets.

## Results

### 2.1 Scaffolding model with using co-barcoding information

Scaffolding is a process to determine the order and orientation of sequences by correlations between an arbitrary pair of sequences, which could come from different linkage information [23]. If there exists a quantitative relation between linkage information and distance, then the distance between two sequences could be estimated too. In stLFR reads, both the PE and co-barcoding information are contained. Only a scaffolding model for the co-barcoding information would be introduced in this section because models for the PE information have been discussed intensively for NGS reads [24-26]. In ideal SLR datasets, reads co-barcoded with the same barcode come from the same LDF. If these co-barcoded reads can be aligned to two contigs, there does exist overlaps between the LDF and both the two contigs at the same time, indicating that there exists an LDF linkage between these two contigs. Because the read coverage depth of an LDF in most of SLR dataset is low, the overlap between an LDF and two contigs is only a necessary condition of sharing barcodes between these two contigs, but not a sufficient and necessary condition. However, considering the unbiased randomness assumption about generations of LDFs from a genome and reads from an LDF, the correlation strength between two contigs could be approximated as the length of the linkage region as shown in Figure 1A, where the start points of LDFs falls. To concisely show the scaffolding model, both LDFs and contigs with constant lengths were used in Figure 1A. The length of a linkage region, where LDFs have overlaps with two contigs at the same time, is equal to difference between the length of an LDF and that of a gap. This indicates that the correlation strength of the LDF linkage between two contigs decrease with increasing the gap size. This is key to order and orientate contigs by using LDF linkages as shown in Figure 1B and 1C.

**Figure 1.**
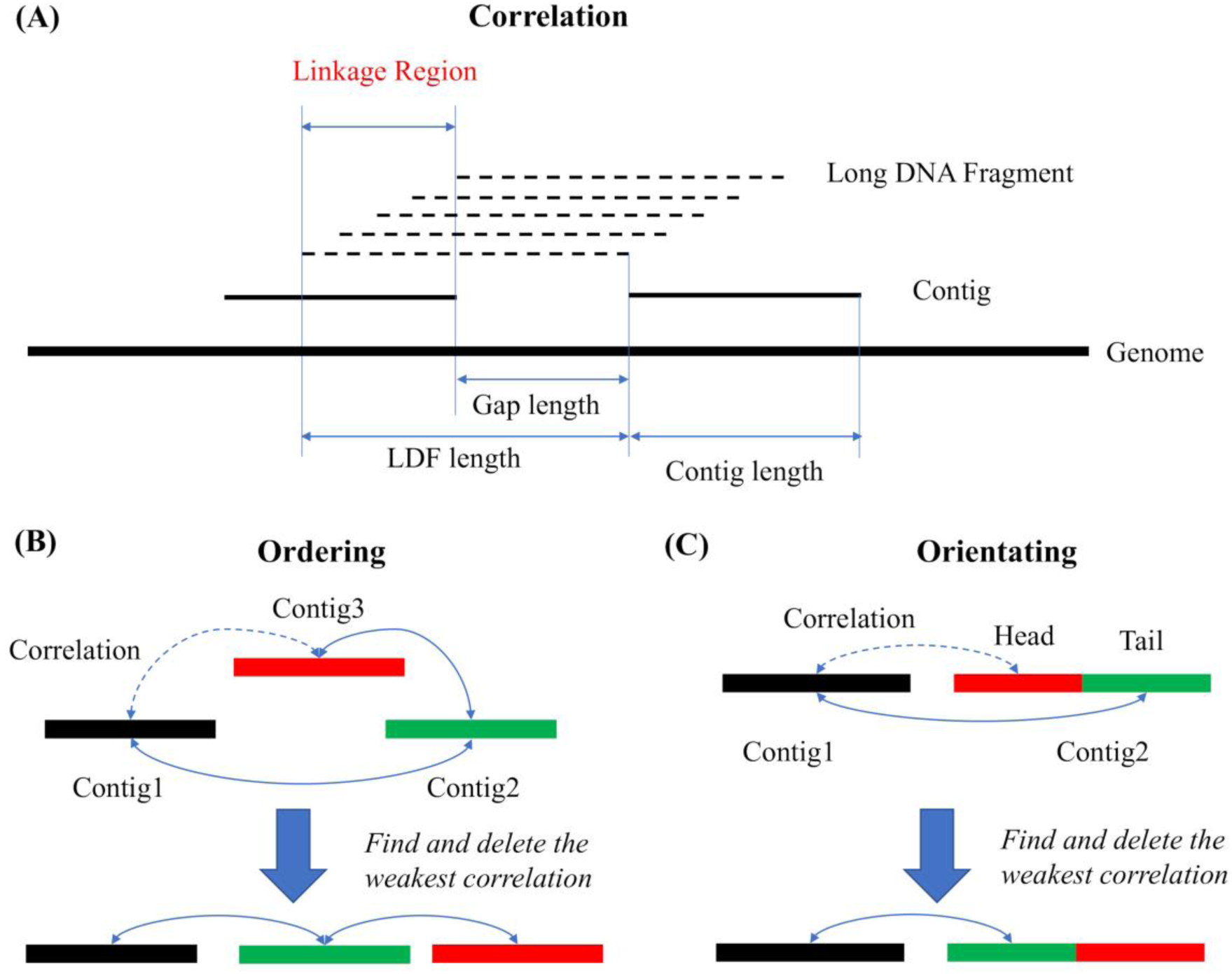
Scaffolding model of the correlation between two contigs, ordering of three contigs and orientating of a given contig by its neighboring contig with using co-barcoding information.

In ordering three contigs with a correct order Contig1, Contig2, Contig3 as shown in Figure 1B, the gap between Contig1 and Contig3 is the largest, which is the sum of the gap between Contig1 and Contig2, gap between Contig2 and Contig3, and length of Contig2. This means the correlation strength between Contig1 and Contig3 would be the weakest one. Thus, if we can find and delete the weakest correlation in the graph as shown in the upper part of Figure 1B, we can determine the order of these three contigs as shown in the lower part of Figure 1B. Since a linkage of an LDF is undirected, the determination of the orientation of a contig relative to its neighboring would be transformed into the determination of the order of three contigs as shown in Figure 1C, where Contig2 would be split into two parts named a head and a tail. Thus, we can see that the determinations of the order and orientation by the co-barcoding information belong to the same process, but the length requirement of ordering can be a half of that of orientating for keeping the same accuracy. It should be noted that complexities introduced by non-uniform distributions of the input contig length, gap size between neighboring contigs, and LDF length, read coverage depth of LDF were ignored in the above description of the scaffolding model.

### 2.2 The characters of stLFR reads

For SLR datasets, the number of LDFs per barcode is an important property for downstream analyses and its ideal value is one. To reduce the number of LDFs per barcode, Danko et al., developed Minerva to deconvolve SLR datasets for metagenomics and their results demonstrated that downstream analyses were improved after deconvolution[27]. To evaluate the number of LDFs per barcode, the distance distribution of neighboring reads with the same barcode from the same LDF and those from different LDFs were analyzed and calculated as shown in Figure 2. Similar to the strategy used to analyze the CPT-seq reads [4], the distances between neighboring reads in the reference with the same barcode were calculated after sorting aligned reads based on their genomic coordinates. There are three typical peaks, and the third peak is expanded in the inset (Figure 2A). The first peak corresponds to gaps between paired reads of PE fragments, and its position is about 251 bp. The second peak corresponds to gaps between neighboring reads from the same LDF, which are named intra-gaps, and its position indicates the typical size of the intra-gap, about 2512 bp. The third peak corresponds to gaps between neighboring reads from different LDFs, named inter-gaps, and the typical size of the inter-gap is about 50 Mb. Compared with the distance distribution of the CPT-seq reads [11], the ratio of the peak value between inter-gaps and intra-gaps for stLFR reads is significantly lower, indicating that the average number of LDFs per barcode in stLFR library is much smaller than that of CPT-seq library. The total number of physical partitions in stLFR library are about 50 million magnetic beads with different barcodes, which is much more than those in CPT-seq library or 10XG-linked reads library [5]. The insert size distribution of stLFR reads represents non-Gaussian (Figure 2B), which is different from that in a standard NGS library. All these results demonstrate that the properties of stLFR reads are a general SLR dataset, and a general scaffolding algorithm is required to efficiently exploit the PE and co-barcoding information of stLFR reads.

**Figure 2.**
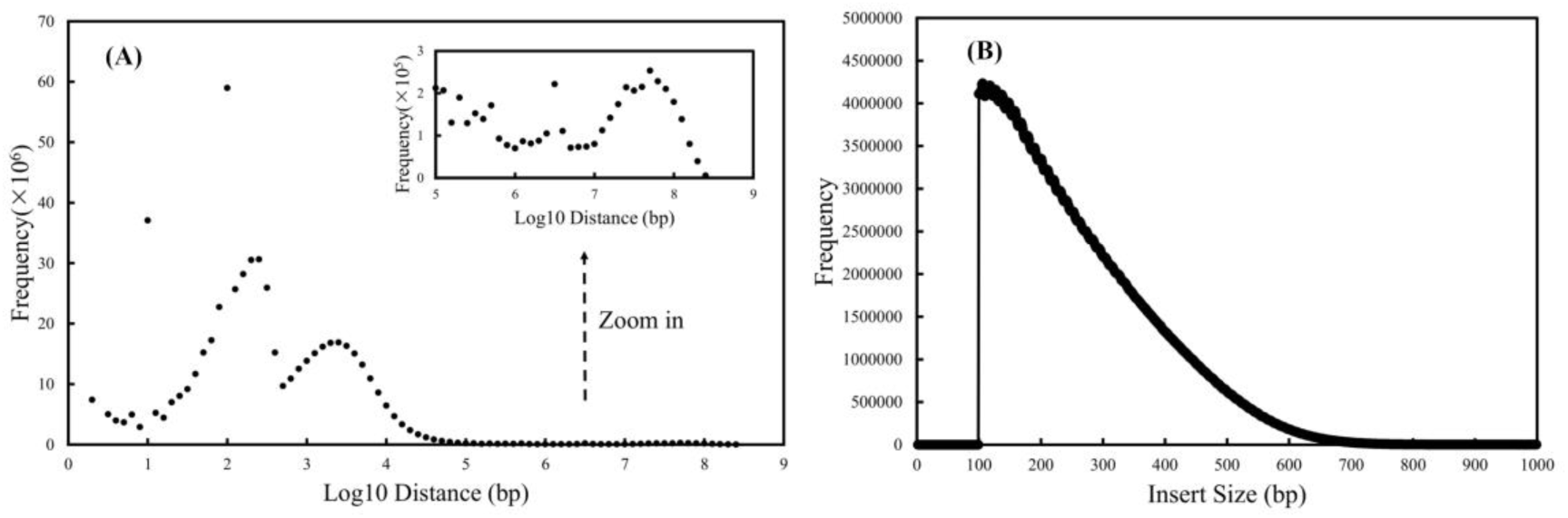
The distribution of distance between neighboring reads with the same barcode (A) and the insert size distribution of read pairs for stLFR reads (B).

According to the scaffolding model, a correlation of the co-barcoding information between contigs could be provided. The number of shared barcodes (NB) between two contigs was used in fragScaff and ARKS, where the fluctuation of LDF coverage depth on the sample genome was ignored. The fluctuation comes from the fact that LDFs are randomly captured in stLFR library. To reduce effects of the fluctuation, JS was used in our work as defined in section 4.1. To understand the advantage of JS relative to NB, both of them between two bins with a fixed size (5000 bp) were analyzed for all pairs of bins with a series of fixed distances (500, 1000, 2500, 5000, 7500, 10000, 20000, 30000 bp) in the chromosome 19 (Chr19) of the human as shown in Figure 3. In Figure 3A and 3B, mean values at different distances are shown, and both JS and NB monotonically decrease with increasing the distance, which is defined as the difference of the leftmost position between two bins. This indicates that these two kinds of correlation are valid to be used to determine the order and orientation of contigs. The property of monotonical decrease of JS comes from co-barcoded reads could also been verified by the dependence of JS on distance for stLFR reads and random barcoded reads as shown in Figure S2. In Figure 3C and 3D, the normalized density distributions of JS and NB at different distances are shown, and both of their distributions are similar to the normal distribution. According to the overlap between two distributions in the figures, we can observe that overlaps for NB are obviously larger than those for JS. The overlap is larger, the probability of making order or orientation errors is larger. The above results demonstrate that JS would be more efficient than NB. The efficiency of JS is also confirmed by the data for cases with bin sizes 1200 and 20000 bp as shown in Figures S3 and S4. It is noted that the efficiency of both JS and NB is dependent on the bin size.

**Figure 3.**
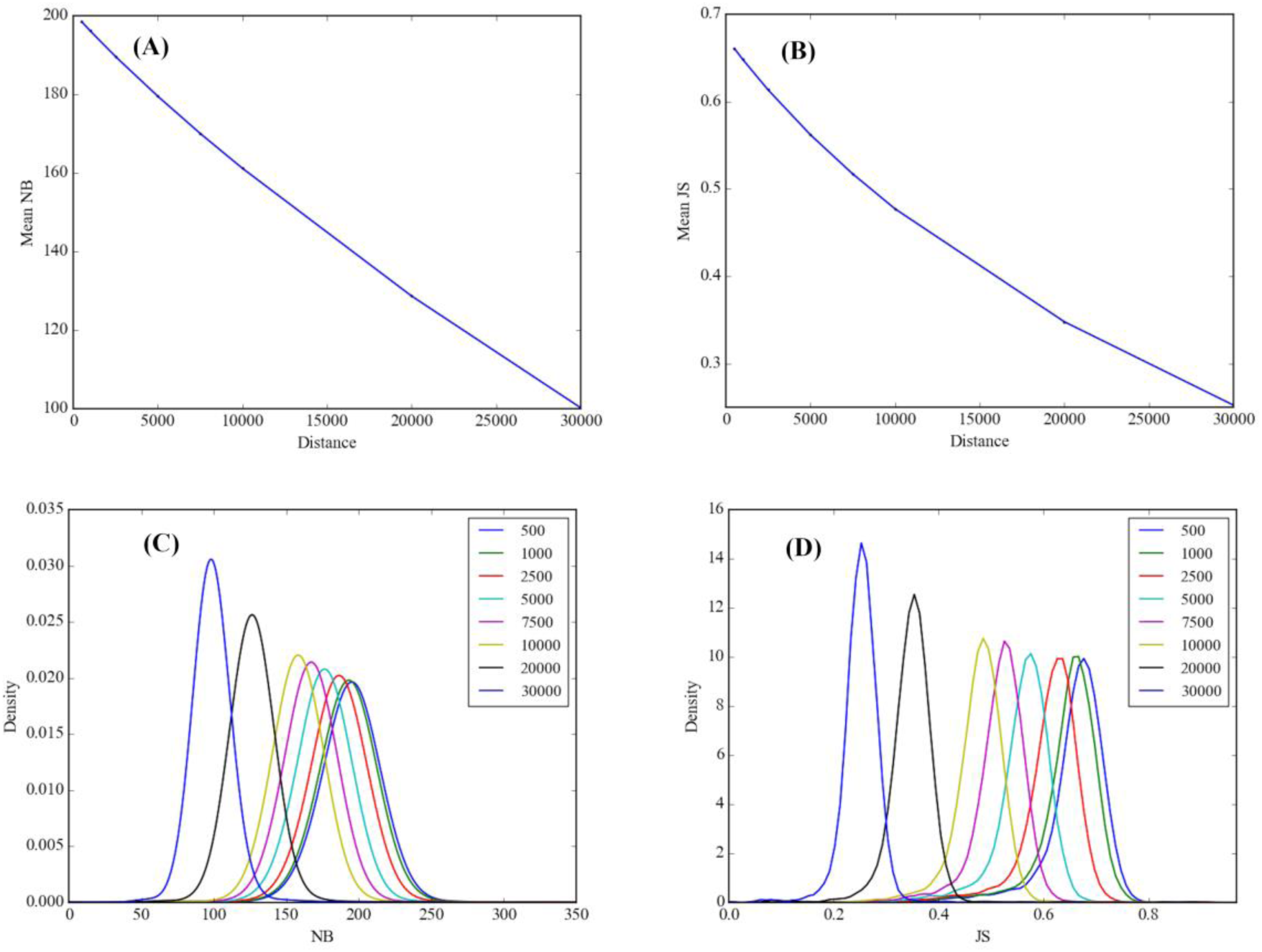
Mean NB of two bins at different distances (A), mean JS of two bins at different distances (B), distribution of NB of two bins at different distances (C), distribution of JS of two bins at different distances (D) with bin size of 5000 bp.

### 2.3 Assembly results with only using stLFR reads

The MaSuRCA contigs of HG001 were obtained by breaking the scaffolds at the positions of unknown bases “N”, where the scaffolds were assembled by MaSuRCA with only using stLFR reads. The evaluations of MaSuRCA contigs are listed in Table S1 and the run parameters of MaSuRCA are listed in Table S4. To evaluate the efficiency of SLR-superscaffolder using the PE and co-barcoding information of stLFR reads, we benchmarked SLR-superscaffolder and other SLR scaffolding tools including fragScaff, Architect and ARKS. For each SLR scaffolder, run parameter sweeps were completed for the dataset of Chr19, and the optimal results are listed in Table S5.

For MaSuRCA contigs, scaffolds assembled by SLR-superscaffolder are with the longest contiguity and the highest accuracy. Compared to the evaluation results of MaSuRCA contigs in Table S1, NG50 of scaffolds is improved by SLR-superscaffolder about 1349-fold from 13.1 kb to 17.6 Mb, and NGA50 is improved about 29-fold from 13.0 kb to 380.5 kb. Except our tool, fragScaff generates scaffolds with the highest quality, where NG50 and NGA50 reach 400.9 kb and 17.5 kb. It is noted that the contiguity and accuracy of scaffolds assembled by fragScaff, Architect and ARKS are significantly lower than those listed in the work of ARKS. One possible reason would be that NG50 of input assemblies in this work is only about 13 kb, which is significantly smaller than that in the work of ARKS about 5 Mb. Compared to scaffolds assembled by MaSuRCA, where co-barcoding information was not used, scaffolds assembled with the co-barcoding information are improved for all SLR scaffolders, especially for SLR-superscaffolder. The above results indicate that the co-barcoding information of stLFR reads can be used to improve draft assembles. It is notable that SLR-superscaffolder can make a greater improvement than other SLR scaffolders in terms of input contigs with relatively short contiguity.

In our top-to-bottom scheme, the ordered scaffolds generated in the ordering step determine the contiguity of final scaffolds and also have direct effects on the accuracy of following steps. To make ordering step with higher accuracy, the screening on an MST was designed to reduce negative effects from non-ideal seed contigs, which include contigs with misassemblies or long repeat sub-sequences. Based on the QUAST evaluation of contigs, the number of non-ideal seed contigs in the MST of MaSuRCA contigs was counted to demonstrate the effects of screening algorithm. In the initial MST, there are 3083 non-ideal seed contigs out of total 182046 contigs. After performing the screening algorithm, there are still 2225 non-ideal seed contigs out of total 179719 contigs. In other words, the ratio of non-ideal seed contigs for the screened contigs is about 0.33, which is larger than that for the initial contig set about 0.02. This indicates that the screening algorithm has high efficiency to find non-ideal seed contigs in an MST, although it cannot completely identify all non-ideal seed contigs.

To clarify effects of the screened contigs on scaffolding, the properties of the MST before and after screening were analyzed as shown in Table 2, where classifications of the nodes and edges in an MST were defined in section 4.7. For edges in the MST before screening, the number of 1-order edges are 172036, which is 96% of total. This result demonstrates that 1-order edges could be efficiently determined by the MST algorithm. The numbers of 2-order and higher-order are only 877 and 613 respectively. The number of error edges is 6441. Dividing the number of edges by the number of non-ideal seed contigs, we can see that the average number of edges of non-ideal seed contigs is about 2.09, which is obviously larger than that of correct contigs. Insight of the numbers of different edges, it is obvious that error edges may have more negative impacts on the MST relative to the higher-order edges. For nodes in the MST before screening, the number of tip junctions and long junctions are 1826 and 354, and those of linear and tip nodes are 177109 and 2758. In our algorithm, the order of neighboring contigs of long junctions cannot be determined by the co-barcoding information. Since there exist 354 long junctions, it is not possible to find a main truck of the MST to scaffold most of contigs as realized in fragScaff. In the MST after screening, the numbers of error edges and higher-order edges are significantly reduced by 2524 and 181, but those of 1-order and 2-order edges are slightly reduced by 20 and 12, compared to the MST before screening. Meanwhile, there are no more long junctions. These results indicate that not only the accuracy of getting 1-order edges is strengthened but also the complexity of the MST is reduced. Whether the contigs of the junction or in the local graphs of the junction are non-ideal seed contig or not was analyzed and the results are listed in Table S7 and S8. From those tables, 70.3% of long junctions were non-ideal seed contigs, and 88.4% of local graph around long junctions contain non-ideal seed contigs. This indicates that long junctions of the MST have a strong correlation with non-ideal seed contigs and also explains why our screening strategy makes sense.

**Table 1.** Evaluation summary of assemblies using MaSuRCA contigs as input assemblies for HG001.

|  | SLR-superscaffolder | fragScaff | Architect | ARKS | MaSuRCA |
| --- | --- | --- | --- | --- | --- |
| Number of scaffolds (>1000bp) | 119,630 | 166,476 | 307,205 | 213,076 | 300,831 |
| Largest scaffold (bp) | 62,433,675 | 4,043,217 | 202,889 | 2,756,390 | 177,746 |
| Total assembled length (bp) | 3,345,341,888 | 3,672,256,418 | 3,038,310,109 | 2,908,519,565 | 2,907,642,015 |
| NG50 (bp) | 17,657,864 | 400,954 | 14,042 | 37,406 | 13,405 |
| NGA50 (bp) | 380,495 | 17,539 | 13,705 | 15,020 | 13,232 |
| Relocation | 11,015 | 92,267 | 3,828 | 51,228 | 1,648 |
| Inversion | 2,939 | 5,349 | 1,637 | 5,190 | 180 |
| Translocation | 2,472 | 2,294 | 903 | 10,828 | 849 |
| Number of misassemblies | 16,426 | 99,910 | 6,368 | 67,246 | 4,475 |

**Table 2.** Statistics of nodes and edges in the MST before and after screening the long junctions.

|  | Tip<br>junction | Long<br>junction | Tip<br>node | Linear<br>node | 1-order<br>edge | 2-order<br>edge | High-order<br>edge | Error<br>edge |
| --- | --- | --- | --- | --- | --- | --- | --- | --- |
| Before screening | 1826 | 354 | 2758 | 177,109 | 172,036 | 877 | 613 | 6441 |
| After screening | 1149 | 0 | 2430 | 176,140 | 172,016 | 865 | 432 | 3917 |

### 2.4 Effects of the length threshold of seed contigs

To analyze effects of different parameters in our tool, the pipeline of combing MaSuRCA and SLR-superscaffolder were conducted with only using stLFR reads of Chr19. Comparing scaffolding results of MaSuRCA to those of SLR-superscaffolder listed in Table S6, both scaffold NG50 and NGA50 values are substantially improved about 316-fold from 27.5 kb to 8.7 Mb and 33-fold from 26.3 kb to 873.7 kb, respectively. These results also demonstrate that both the PE and co-barcoding information is used with high efficiency and precision by SLR-superscaffolder.

The contiguity and accuracy of input contigs have strong effects on the quality of the assembly but the accuracy could not be determined without a reference for *de novo* assembly. Thus, we concentrated on evaluating effects of contiguity by changing the length threshold to choose seed contigs. As shown in Figure 4, with increasing length threshold, scaffold NG50 monotonically decreases, while NGA50 reaches a saturation peak between 5 kb and 10 kb. In terms of major misassemblies, the numbers of inversion and relocation errors monotonically decrease with increasing the length threshold, and the decreasing rate in the region of small thresholds is obviously higher than that in the region of large thresholds. The dependence of the length threshold on scaffold NG50 demonstrates that the connectivity of co-barcoding scaffold graph can be enhanced by involving more contigs when using a small threshold. However, shorter contigs are more inclined to introduce misassemblies. Thus, to get an optimal assembly by the co-barcoding information, it is very important to make a good balance between the connectivity and complexity of a co-barcoding scaffold graph through tuning the number of short contigs. Although the balance is not determined only by the length threshold of input contigs, the saturation peak of scaffold NGA50 indicates that our tool is robust to achieve a relatively optimal balance.

**Figure 4.**
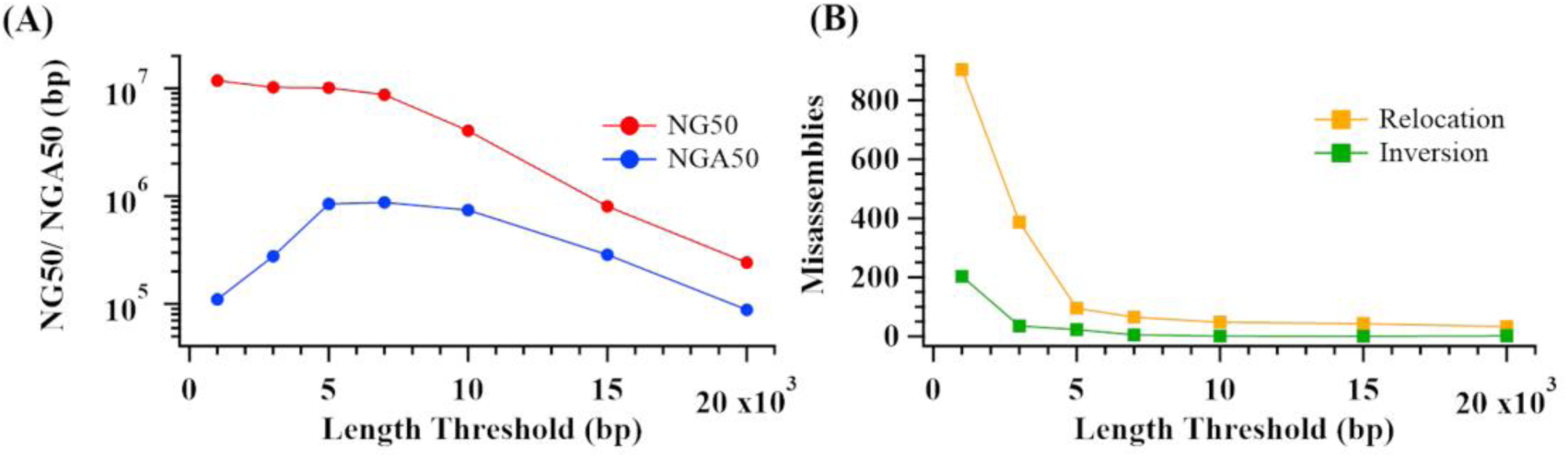
Quality of scaffolds assembled by SLR-superscaffolder with different length thresholds of seed contigs.

In addition, effects of local scaffolding by PE information were tested using dataset of Chr19, and the results are listed in Table S6. Comparing results with local scaffolding to those without, although improvements of both the contiguity and accuracy are not substantial, the decreasing ratio of inversion errors is as high as about 2.5-fold, indicating that our top-to-bottom scheme is an efficient way to make good use of the complementary between the PE and co-barcoding information.

### 2.5 Assembly results by combining stLFR reads with other sequencing reads

As a standalone scaffolding tool, SLR-superscaffolder can be easily implanted in a hybrid assembly strategy, where stLFR reads and other kinds of sequencing datasets could be used together. In this work, two cases were tested for all SLR scaffolders: one is the combination of stLFR reads and PCR-free NGS reads, and the other is the combination of stLFR reads and ONT reads. In the first case, the input assembly (SOAP*denovo* scaffold) is consist of scaffolds assembled by SOAP*denovo*2 with stLFR reads and PCR-free NGS reads. In the second case, the input assembly (ONT contigs) is consist of contigs assembled by Canu with ONT reads. The evaluations are listed in Table S1. For both cases, the evaluations of final scaffolds assembled by different SLR scaffolders are listed in Table 3 and the run parameters for each scaffolder are listed in Table S4.

**Table 3.** Evaluation summary of assemblies using SOAP*denovo* scaffolds and ONT contigs as input for HG001.

|  | SLR-superscaffolder | fragScaff | Architect | ARKS |
| --- | --- | --- | --- | --- |
| Human (SOAPdenovo scaffolds) |  |  |  |  |
| Number of scaffolds (>1000bp) | 48,278 | 54,193 | 79,435 | 80,400 |
| Largest scaffold (bp) | 35,605,665 | 59,127,605 | 796,758 | 10,897,000 |
| Total assembled length (bp) | 3,115,923,941 | 3,094,665,921 | 2,659,717,897 | 2,713,878,926 |
| NG50 (bp) | 9,113,260 | 2,346,521 | 54,245 | 468,461 |
| NGA50 (bp) | 1,510,911 | 101,813 | 44,836 | 59,899 |
| Relocation | 2,373 | 32,588 | 1,104 | 20,906 |
| Inversion | 105 | 1,866 | 39 | 110 |
| Translocation | 3,694 | 2,926 | 875 | 4,060 |
| Number of misassemblies | 6,172 | 37,380 | 2,018 | 25,076 |
| Human (ONT contigs) |  |  |  |  |
| Number of scaffolds (>1000bp) | 807 | 1,051 | 1,474 | 876 |
| Largest scaffold (bp) | 90,148,984 | 109,245,684 | 45,826,758 | 170,045,596 |
| Total assembled length (bp) | 2,829,830,390 | 2,828,106,943 | 2,823,722,824 | 2,823,836,148 |
| NG50 (bp) | 21,779,983 | 26,579,775 | 8,806,572 | 39,604,458 |
| NGA50 (bp) | 1,578,910 | 1,592,388 | 1,481,592 | 1,574,345 |
| Relocation | 4,378 | 4,182 | 3,962 | 4,266 |
| Inversion | 64 | 63 | 60 | 61 |
| Translocation | 1,575 | 1,453 | 1,383 | 1,607 |
| Number of misassemblies | 6,017 | 5,698 | 5,405 | 5,934 |

For SOAP*denovo* scaffolds, scaffolds assembled by SLR-superscaffolder are with the longest contiguity and the highest accuracy. Compared to evaluation results of SOAP*denovo* scaffolds in Table S1, NG50 of scaffolds assembled by SLR-superscaffolder is improved about 227-fold from 40.1 kb to 9.1 Mb, and NGA50 is improved about 44-fold from 34.3 kb to 1.5 Mb. Except our tool, fragScaff generates scaffolds with the highest quality, where NG50 and NGA50 reach 2.3 Mb and 101.8 kb. As discussed for results of MaSuRCA contigs, NG50 of SOAP*denovo* scaffolds is much shorter than that of those used in the work of ARKS. For ONT contigs, all scaffolders can make an obvious improvement of contiguity but little improvement of accuracy. Compared to the evaluations of ONT contigs in Table S1, SLR-superscaffolder produces an improvement of contiguity about 3.3-fold from 6.6 Mb to 21.8 Mb, which is slightly lower than that by ARKS (about 6-fold) and fragScaff (about 4-fold). The largest improvement of NGA50 is produced by fragScaff about 1.13 from 1.4M to 1.6M, and the NGA50 obtained by our tool is very close to that of fragScaff. As shown in the evaluation of ONT contigs in Table S1, although the quality of ONT contigs is high, the average number of misassemblies of an ONT contig is about 3.2. Considering that contigs with misassemblies are more likely to be screened in our screening algorithm, the connectivity of the co-barcoding scaffold graph composited of long ONT contigs would be significantly reduced. The above results indicate that the accuracy of an input assembly is very important to the accuracy of the final assembly for all SLR scaffolders but SLR-superscaffolder is more robust to the contiguity of input assembly.

### 2.6 Overall performance

In addition, we evaluated the running time of each scaffolder for three inputs on the same computational platform (Intel(R) Xeon(R) CPU E7-4890 v2 2.80GHz, 60 core, 120 threads and 3 T RAM in total) as shown in Figure 5. All computations were limited to 20 threads. The comparison showed that ARKS has the best overall performance because it adopted a *k*-mer-based mapping strategy to avoid time-consuming pairwise aligning. In spite of the alignment-based gain of the co-barcoding information, SLR-superscaffolder run approximately 1.5-, and 4.3-fold faster than fragScaff and Architect on average, respectively. The running time statistics for each step of SLR-superscaffolder as shown in Table S9 demonstrate that the data preparing step including stLFR read mapping and co-barcoding information assignment is the most time-consuming, averaging 58.3% of the total. Note that we did not compare the peak memory consumption since the maximal usage depended on the aligners instead of the scaffolders themselves.

**Figure 5.**
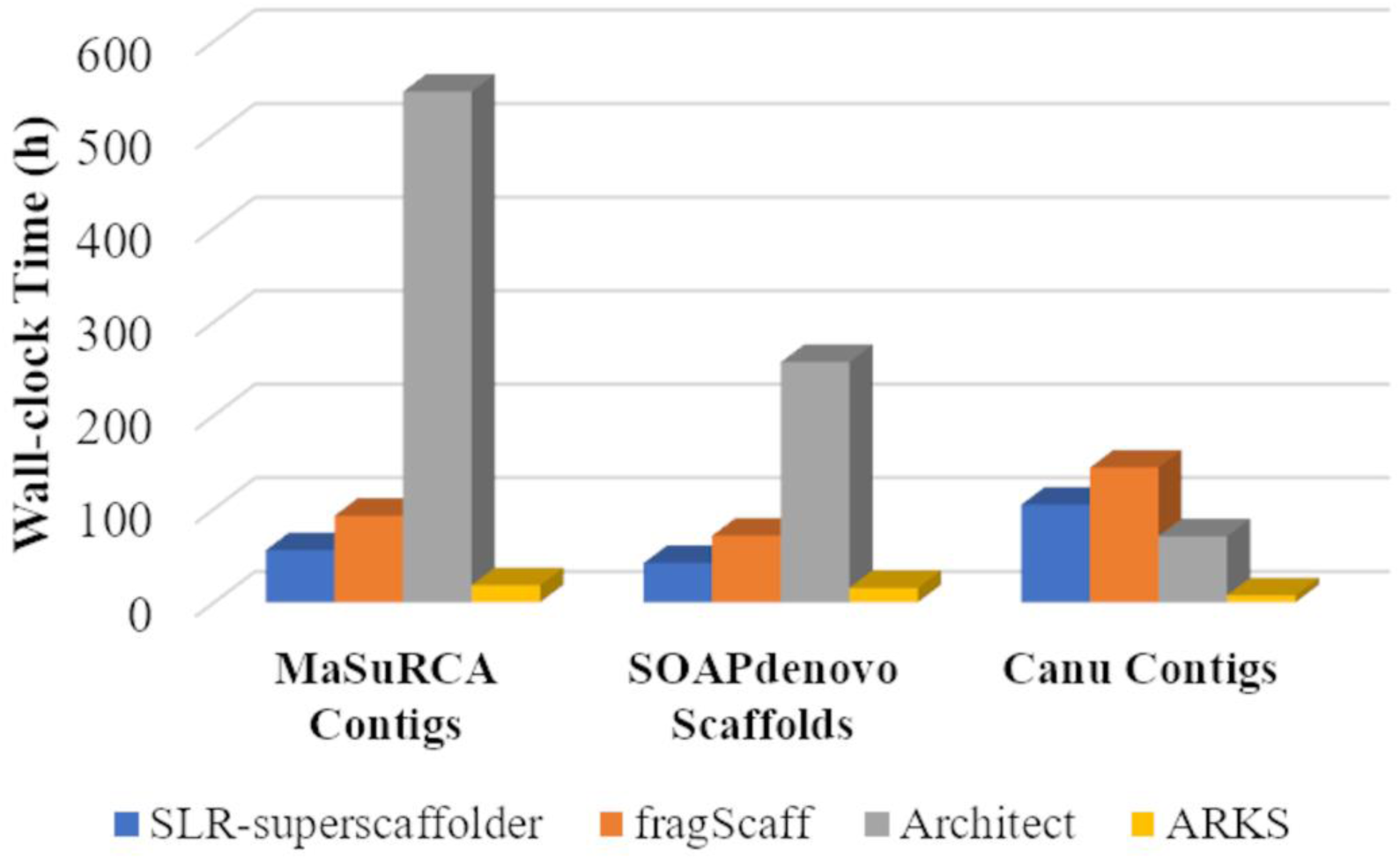
Histograms of time consumption of four scaffolders (SLR-superscaffolder, fragScaff, Architect, ARKS) for three input assemblies (MaSuRCA contigs, SOAP*denovo* scaffolds, Canu contigs).

## Discussion

According to statistical properties of stLFR dataset by analyzing stLFR reads on the reference, the statistical properties of the PE and co-barcoding information are exhibited and the results demonstrate that a stLFR dataset is a general SLR dataset with irregular insert size of PE fragments and a fewer number of LDFs per barcode. To use the complementary information with high efficiency in *de novo* genome assembly, we introduced a new top-to-bottom scaffolding algorithm in SLR-superscaffolder according to the scaffolding model of the co-barcoding information proposed in this work. In our top-to-bottom scheme, the co-barcoding information with longer correlation length is determined prior to the PE information and the ordering step with lower requirement of input contig length is determined prior to the orientating step. In our tests of the human whole genome, SLR-superscaffolder produces about several hundred-folds improvements of scaffold NG50 with high accuracy relative to input assemblies for input assemblies assembled by NGS reads. These results demonstrate that the co-barcoding information from stLFR library can be used to improve the quality of draft genomes assembled with different sequencing datasets.

The SLR-superscaffolder is the first SLR scaffolder that considers to reduce effects of misassemblied contigs in input assemblies and provides systematical screening of these contigs on the scaffold graph by combining with the topology of junctions in the maximum-weight MST. Our results show that the screened seed contigs have strong correlation with non-ideal seed contigs. Compared with other SLR scaffolders, SLR-superscaffolder produces scaffolds with longer contiguity and higher accuracy for different input assemblies, indicating that our tool is with highly efficient and robust to use the co-barcoding information of stLFR reads. It is important to note that all other SLR scaffolders are specifically designed for their corresponding SLR library dataset other than stLFR reads and the control parameters of each SLR scaffolder used in different datasets may not be optimal although parameter sweeps have been conducted for the stLFR dataset of Chr19.

As a standalone scaffolder, SLR-superscaffolder improves the quality of assembly results from other types of library (such as standard NGS or SMRT libraries) using the co-barcoding information in stLFR reads. Although SLR-superscaffolder is initially designed for stLFR reads, the co-barcoding information in other types of SLR datasets could also been exploited with an appropriate format conversion considering the general properties of stLFR reads. Furthermore, since our approach is highly modularized, each step in SLR-superscaffolder will be separately improved when combined with other types of sequencing datasets such as SMRT or mate-pair in our future work.

## Methods

### 4.1 Jaccard similarity

In practice, a function of shared barcodes between two sequences should be used to represent the correlation strength. In this work, JS was selected by comparing different quantities as discussed in section 2.2. To avoid effects of the contig length variation, JS between contig *m* and contig *n* was defined as the maximal JS between paired bins with the same size from each two contigs as following,

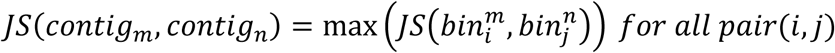

where 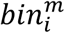 was the *i*_*th*_ bin in contig *m*. JS between bins from different contigs was calculated as following,

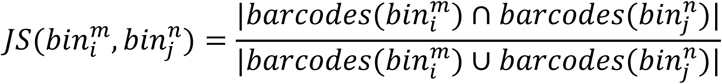

where *barcodes* 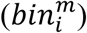 was the set of barcodes whose corresponding reads were aligned to the 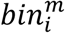. The bins used in this work were chopped without any gaps or overlaps along a contig.

### 4.2 Overview of the algorithm and data preparing

In our tool, both the PE and co-barcoding information were used to scaffold a draft assembly. In our strategy, the co-barcoding information with large correlation length was used firstly to scaffold contigs with long length on a global scale and then the PE information was used to scaffold contigs with short length on a local scale. In scaffolding with the co-barcoding information, the determination of the order and orientation were divided into two modules, where the order was determined firstly, then the orientation. Finally, the gap size between neighboring contigs was estimated. Overall, five modules were integrated orderly as shown in Figure S1, including data preparing, ordering, orientating, local scaffolding, estimating gap size. Because the co-barcoding information with longer correlation length and ordering with a lower requirement of the input contig length were determine firstly as above, the strategy adopted in our tool is called a top-to-bottom scheme.

For SLR-superscaffolder, there are two kinds of input including a SLR dataset and a draft assembly. A draft assembly can be contigs or scaffolds assembled with any types of datasets, but only contigs were used in the description of our methods. Before scaffolding, two kinds of data preparing were conducted. One is to get the relation between reads and contigs to derive both the PE and co-barcoding correlation between contigs. In this preparing, BWA (version 0.7.17) [28] was used to align stLFR reads to contigs, and then the left most positions of alignments of reads were used to define the relation between reads and contigs. The other is to choose a subset of contigs to be scaffolded by the co-barcoding information in our top-to-bottom scheme. To reduce complexities caused by repeat contigs and the low resolution of the co-barcoding information, long contigs which appear only once in a genome were chosen as seed contigs in scaffolding with using the co-barcoding information. In practice, seed contigs were determined by those whoes length is longer than a threshold (default 7000 bp) and with coverage depth around the average value from 0.5 to 1.5 times. Because the distribution of coverage depth is not uniform and misassemblies would be contained in input contigs, there would exist non-ideal seed contigs obtained in this step. Therefore, it is necessary to reduce negative effects of these non-ideal seed contigs on scaffolding.

### 4.3 Ordering

In our top-to-bottom scheme, the order between contigs at global scale was firstly determined by using the co-barcoding information. The contiguity of scaffolds after this step determines the contiguity of final scaffolds and the accuracy of them is important to the accuracy of following steps. For keeping the contiguity and accuracy in the ordering step, the screening of non-ideal seed contigs was firstly conducted as shown in Algorithm 1. In our algorithm, the graph-based algorithms and topology of a tree graph were used as described in section 2.1. A co-barcoding scaffold graph is a weighted undirected graph. The nodes are seed contigs chosen in the data preparing step. The weighted edge between two nodes exists only when JS between two contigs is larger than a threshold and the weight is the value of JS. Junctions and branches were focused on in the use of topological properties of a tree graph. The junction is a node with a degree higher than two. The branch is a linear path from a tip node, whose degree is one, to the nearest junction of it. If the number of nodes in a branch is less than three, the branch was defined as a tip branch, otherwise defined as a long branch. According to the length of branches, the junction connected with more than two long branches was defined as a long junction, otherwise a tip junction. The maximum-weight MST of a co-barcoding scaffold graph was obtained by using the Prim’s algorithm. If a MST contains many long junctions, it was not able to efficiently order contigs by extracting the trunk of the MST as conducted in fragScaff [11]. By analyzing properties of contig around junctions in the MST of the MaSuRCA contigs as shown in Table S7 and S8, we found long junctions have strong correlation with the non-ideal seed contigs. In the while loop of the screening algorithm, the end condition was set by the number of iterations and ratio of screened contigs to all contigs to avoid a significant reduction of connectivity in the co-barcoding scaffold graph. The default number of iterations was 100, and ratio of screened contigs was 0.05%. It is noted that only a part contigs were ordered in this step, where contigs in tip branches and long junctions were ignored.

#### Algorithm 1: Ordering

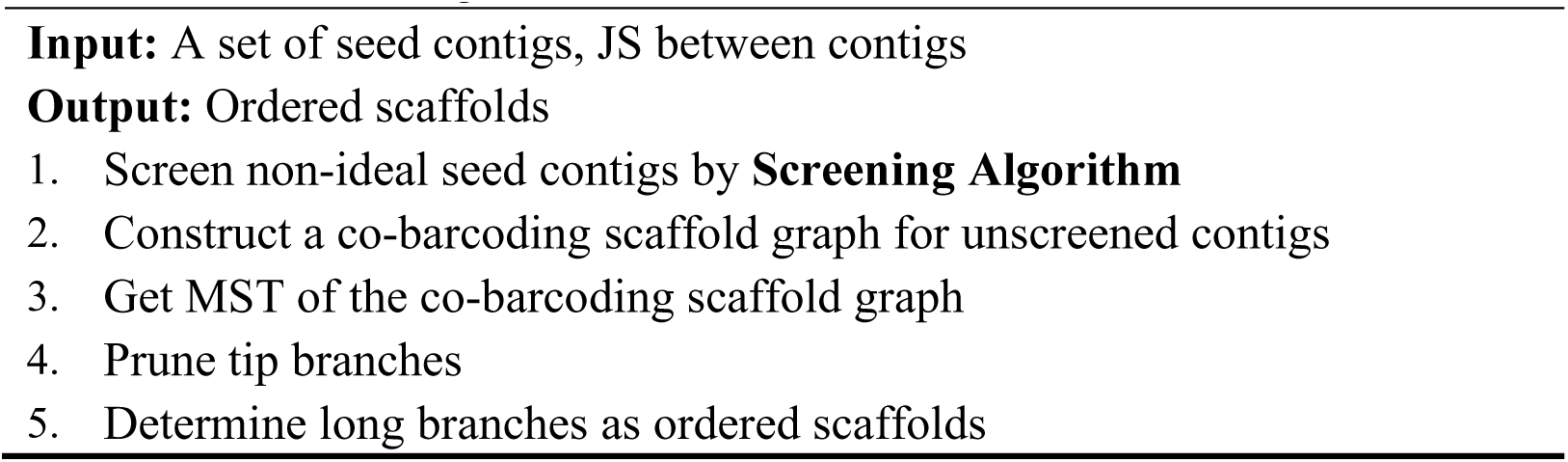

#### Algorithm 2: Screening

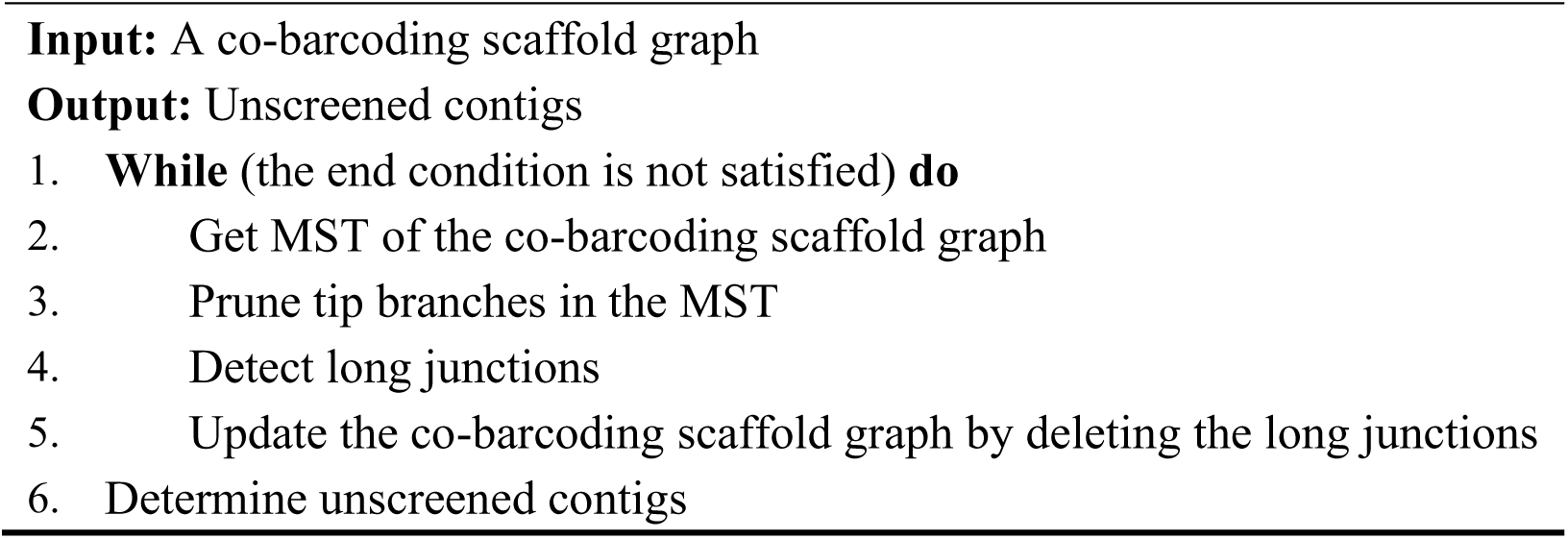

### 4.4 Orientating

Because the co-barcoding information is undirected, contigs would be divided into two parts at the middle point of a contig as shown in Figure 1C for orientating with a similar strategy that has been also adopted in other tools. In this work, the head of a contig is the part from 5’ terminal to the middle point, and the tail is the part from the middle point to 3’ terminal. However, different from other tools, where the order and orientation were determined at the same time, the information of neighbor contigs in an ordered scaffold could be used to determine the orientation of each contig in our strategy. To make use of this local information in an ordered scaffold, we designed a consensus strategy as shown in Algorithm 3 to determine the orientation of each contig. In this strategy, one neighboring contig may provide one support for an orientation state of a contig as shown in Figure 1C. For a given contig, a contig in the same ordered scaffold with difference of order relative to it less than a threshold was defined as its neighboring contig. The threshold of the neighboring contig was 4. The orientation of a contig has two states: one is up meaning that the direction from 5’ terminal to 3’ terminal of the contig is the same with that of the ordered scaffold, the other is down meaning that the direction is opposite. A support of a neighboring contig is determined by the relative magnitude of JS between the head and the neighboring contig (JS_Head) and JS between the tail and the neighboring contigs (JS_Tail). The JS_Head and JS_Tail were also derived from the alignment relations between barcodes and contigs obtained in the data preparing step. To increase computational efficiency to determine orientation states for a paired neighboring contigs in one calculation, neighboring contigs were divided into two parts practically.

#### Algorithm 3: Orientating

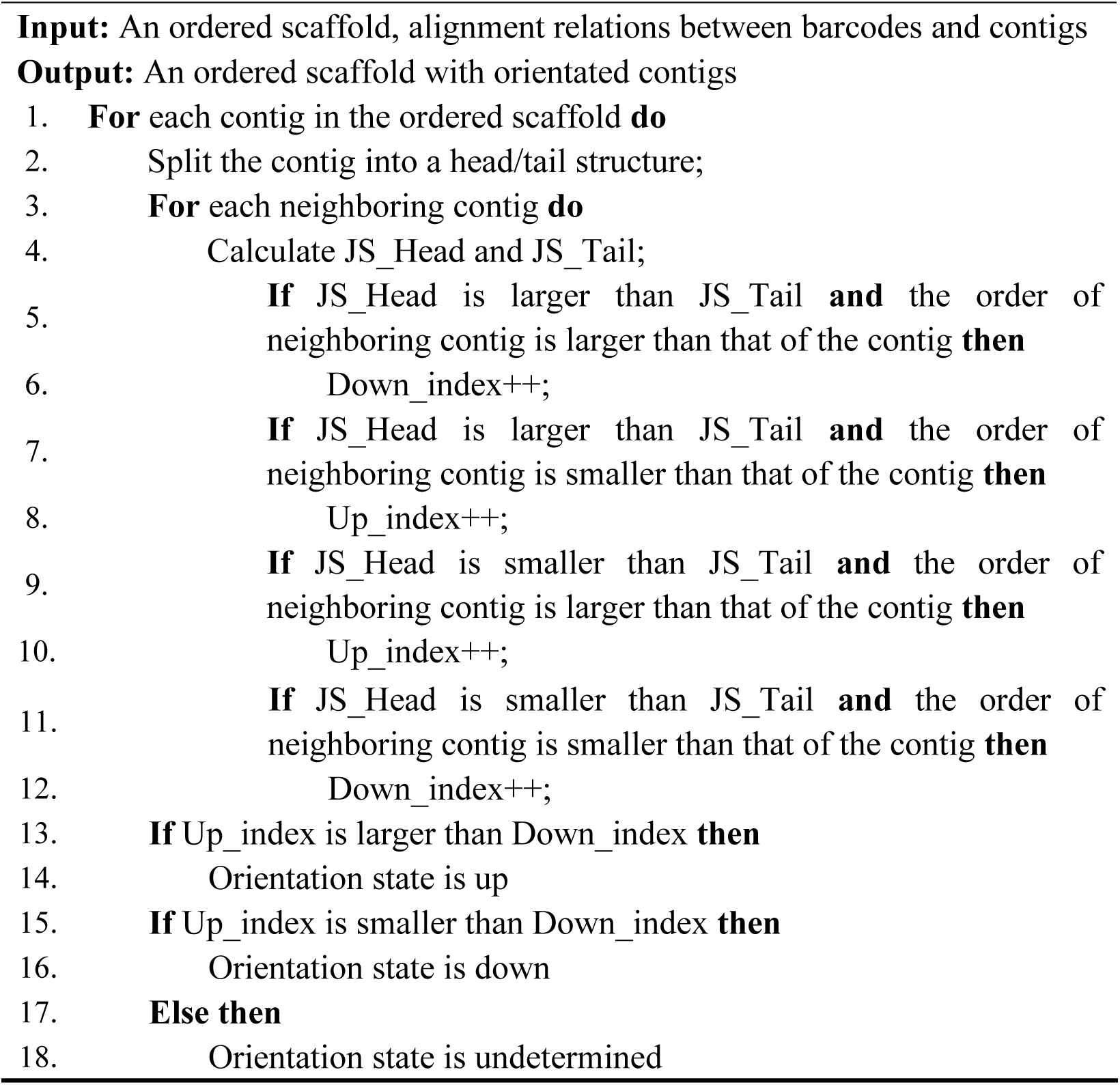

### 4.5 Local scaffolding

In above steps, the PE information of stLFR reads was not used and only most of seed contigs were ordered and oriented by the co-barcoding information. The unscaffolded contigs include non-seed contigs labeled in data preparing, those of tip branches in the maximum-weight MST, and those screened in the ordering step. In this step, we inserted the first two types of unscaffolded contigs into oriented scaffolds by combining the PE with the co-barcoding information for paired neighboring contigs of each gap one by one with Algorithm 4. For paired contigs of a gap in scaffolds after orientating, unscaffolded contigs were clustered as candidates for local scaffolding when the number of shared barcodes to paired contigs of a gap was larger than a threshold. The threshold in the clustering was 10. In the local directed PE scaffold graph, the nodes are consisted of candidate contigs and the paired contigs of the gap, and the directed edges refer to a connection between two contigs, where the number verified by the PE information is larger than a threshold, which was 3 practically. The shortest connected path between the paired contigs is determined by using the depth-first search (DFS) algorithm on a directed graph. The above process was similar to the scaffolding process of other NGS assemblers, but our local scaffolding made it more efficient to deal with the complex structures caused by repeat sequences on global scale.

#### Algorithm 4: Local scaffolding

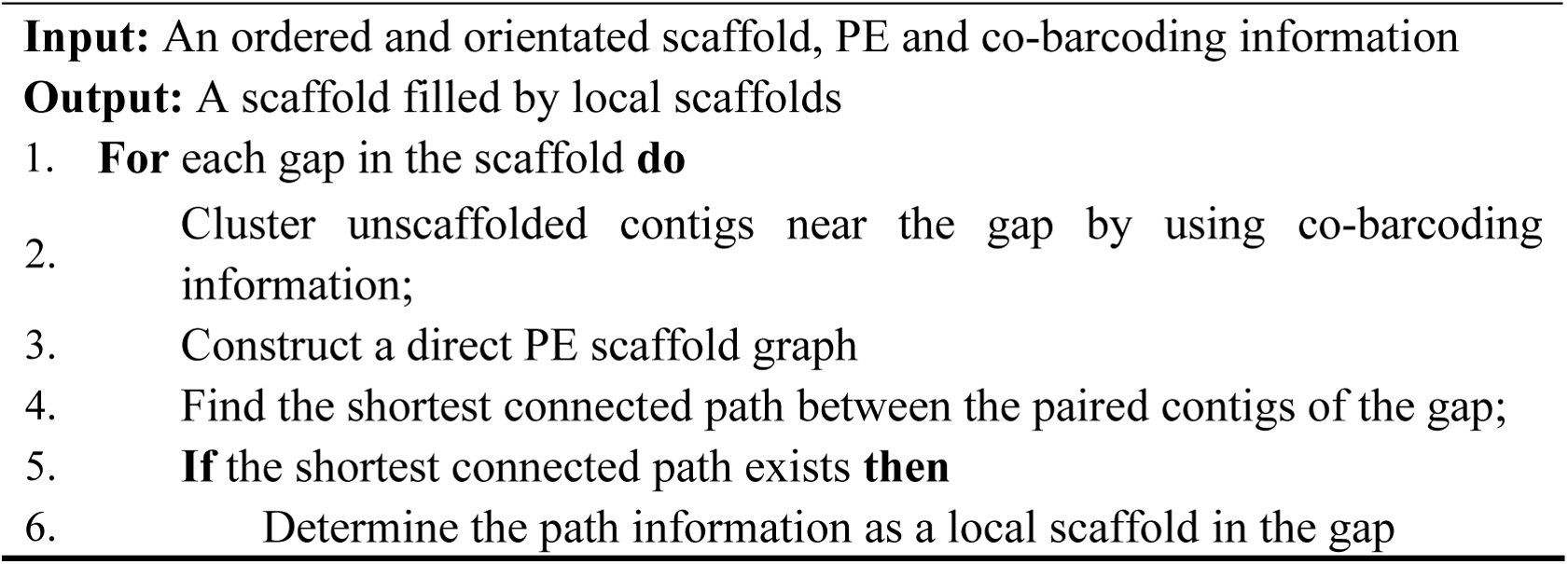

### 4.6 Estimating gap size

Considering that gap sizes between neighboring contigs in scaffolds are useful for further analysis, we estimated them using an empirical relation between distance and JS for gaps constructed by the co-barcoding information (Figure S1E). A similar strategy has been adopted in ARKS [15], where the distance corresponding to a JS was defined as the median value. For gaps constructed by the PE information, the size was uniformly set to 11 bp. Although distances between each two reads with the same barcode were unknown, a rough relation between JS and the distance of two sequences was available according to the statistics in the reference as shown in Figure S2. In practice, the relation was derived from the statistics between sequences on long contigs of input assembly, with a linear fit using the least square method. Finally, gap sizes between contigs were estimated by JS.

### 4.7 Evaluation

The standard metrics of QUAST (version 5.0.2) [29] were used to evaluate the accuracy of assembled results, where the effects of gap sizes were also considered. The correct positions of sequences were determined by the aligning stLFR reads to the reference using Minimap2 (version 2.17-r954-dirty) [30]. Comparing the relative positions of two consecutive sequences on an assembly to those on the reference, all misassemblies were determined. A major misassembly was defined as an alignment difference larger than 1 kb relative to the reference. Major misassemblies were further categorized as three types: relocation, inversion and translocation. Based on different types of misassembly, we could roughly evaluate the accuracy of independent steps in our algorithm and compare with other scaffolding tools. The evaluations of QUAST were run with the parameters (*-m 1000, -x 1000, -scaffolds, -scaffold-gap-max-size 10,000*).

The topological properties of an MST were changed by screening nodes in the ordering algorithm as described above. To completely analyze effects of the screening algorithm on the MST, the edges of the MST were also evaluated by combing the QUAST evaluation results and alignments of scaffolds to the reference. The edges were categorized into four classes including 1-order, 2-order, high-order and error edges. If there does not exist any other contigs between two contigs in the reference genome according to alignments, a correlation between the two contigs was defined as a 1-order edge. If the number of contigs between two contig is equal to one, a correlation between the two contigs was defined as a 2-order edge. If the number is larger than one, the correlation was defined as a high-order edge. For above edges, both two contigs must be without misassembly according to QUAST evaluation. If there exist misassemblies in two contigs, a correlation between the two contigs was defined as an error edge.

### 4.8 Draft assemblies and datasets

In this work, three draft assemblies of HG001 were used as input, including the contigs assembled by MaSuRCA [31] with stLFR reads only (MaSuRCA contigs), the scaffolds assembled by SOAP*denovo2* [32] with stLFR reads and an additional 20-fold PE PCR-free NGS dataset (SOAP*denovo* scaffolds) and the contigs assembled by Canu[33] with about 30-fold Oxford Nanopore technology (ONT) reads from (ONT contigs) Jain et al. work[34]. All evaluations for these input assemblies are listed in Table S1. Except ONT dataset, reads for both stLFR and PCR-free NGS dataset were sequenced by ourselves and have been deposited in the CNSA (https://db.cngb.org/cnsa/) of CNGBdb. The access information of these datasets is listed in Table S2. The detailed information of stLFR and PCR-free NGS reads is listed in Table S3. The stLFR library was constructed by MGIEasy stLFR Library Prep Kit and sequenced by BGISEQ-500. In constructing stLFR library, LDFs longer than 50 kb were produced by shearing input DNA, and randomly trapped into barcoded magnetic beads in a single tube, and then fragmented into short sequences with the same barcode by two transposons [1]. The 20-fold pair-end NGS dataset with the insert size about 390 bp were randomly extracted from the dataset of a PCR-free library, which was constructed by MGIEasy FS PCR-Free DNA Library Prep Set V1.0 (MGI, cat. No. 1000013455) and sequenced by MGISEQ-2000 PE150. For parameter sweeps and data analysis, the stLFR reads of Chr19 were extracted and used according to read alignments of HG001 to the reference.

## Additional files

**Supplementary information** contains the following information:

**Table S1**. Summary of the input assemblies in this work

**Table S2**. Genomics dataset sources

**Table S3**. Summary of stLFR and NGS datasets used in this work

**Table S4**. Control parameters used in different scaffolders for different datasets.

**Table S5**. Evaluation of Chr19 assemblies based on MaSuRCA contigs by different scaffolders with the optimal parameters after the parameter sweeps.

**Table S6**. Evaluation of Chr19 assemblies for different tests.

**Table S7**. Statistics of tip and long junctions before and after conducting the screening algorithm.

**Table S8**. Statistics of local properties of tip and long junctions before and after conducting the screening algorithm.

**Table S9**. Runtime statistics for SLR-superscaffolder step by step.

**Figure S1**. Overall scheme of SLR-superscaffolder.

**Figure S2**. Relations between Jaccard similarity of barcodes and distance for two sequences in the reference for stLFR reads and randomly barcoded reads.

**Figure S3**. Mean NB of two bins at different distances (A), mean JS of two bins at different distances (B), distribution of NB of two bins at different distances (C), distribution of JS of two bins at different distances (D) with bin size of 1200 bp.

**Figure S4**. Mean NB of two bins at different distances (A), mean JS of two bins at different distances (B), distribution of NB of two bins at different distances (C), distribution of JS of two bins at different distances (D) with bin size of 20000 bp.

## Abbreviations

SLR: Synthetic long reads
stLFR: Single tube long fragment reads
NGS: next generation sequencing
LDF: long DNA fragment
CPT-seq: contiguity preserving transposition sequencing
MST: minimum spanning tree
PE: paired-end
10XG-linked reads: 10X Genomics Chromium technology
SMRT: single molecular real time sequencing
HG001: the human whole genome for cell line NA12878
JS: Jaccard Similarity
NB: number of shared barcodes
Chr19: hromosome 19.

## Declarations

### Ethics approval and consent to participate

Not applicable.

### Consent for publication

Not applicable.

### Availability of data and materials

The source codes and instructions of SLR-superscaffolder are freely available on GitHub (https://github.com/BGI-Qingdao/SLR-superscaffolder, licensed under GNU General Public License V3.0). The stLFR dataset of HG001 is available on CNSA of CNGBdb (Access ID CNP0000066). The PCR-free NGS dataset of HG001 is available on CNSA of CNGBdb (Access ID CNP0000602). The ONT Canu assembly of HG001 is available at: ftp://ftp.ncbi.nlm.nih.gov/genomes/all/GCA/900/232/925/GCA_900232925.2_NA127_878-rel5/GCA_900232925.2_NA127878-rel5_genomic.fna.gz.

### Authors’ contributions

LD, GF, and XX contributed to the software design. LD, LG, and MX contributed to the software implementation, and data analyses. WW, SG, XZ, FC and OW contribute to the data curation, collection. XL, LD, and MX contributed to the benchmarking design. All authors contributed to the manuscript writing. LD and XL supervised the project. All authors read and approved the final manuscript.

## Acknowledgements

We would thank Hongmei Zhu, Yinlong Xie and many other BGI-Shenzhen employees for fruitful discussions in the development of SLR-superscaffolder.

