## Supplemental file for "SLR-superscaffolder: a *de novo* scaffolding tool for synthetic long reads using a top-to-bottom scheme"

Table S1. Summary of the input assemblies in this work

| | Datasets | Number of scaffolds (>1000 bp) | Largest scaffold | Total assembled length | NG50 (bp) | NGA50 (bp) | Relocation | Inversion | Number of misassemblies |
|---|---|---|---|---|---|---|---|---|---|
| Chr19 | MasuRCA contigs | 4,083 | 148,918 | 57,024,748 | 25,956 | 25,384 | 114 | 4 | 118 |
| Human whole genome | MaSuRCA contigs | 308,823 | 160,048 | 2,905,719,951 | 13,087 | 12,992 | 1,648 | 180 | 2,677 |
| | SOAPdenovo2 scaffolds | 122,465 | 528,666 | 2,862,817,877 | 40,154 | 34,369 | 982 | 28 | 2,745 |
| | ONT contigs | 1,685 | 38,914,416 | 2,823,828,058 | 6,593,913 | 1,403,850 | 3,944 | 60 | 5,364 |

*Note*: MaSuRCA run with parameters *GRAPH_KMER_SIZE = 63, cgwErrorRate=0.15*. SOAP*denovo2* run with parameters *-K 49, -p 16, -a 700, -d 1*.

Table S2. Genomics dataset sources

| Species | Datatype | Source |
|---|---|---|
| *H. sapiens* NA12878 | stLFR Dataset | ftp://ftp.cngb.org/pub/CNSA/CNP0000066/CNS0007594/CNX0005851/CNR0006062 |
| *H. sapiens* NA12878 | PCR-free NGS Dataset | ftp://ftp.cngb.org/pub/CNSA/CNP0000602/CNS0106271/CNX0086627/CNR0106776 |
| *H. sapiens* NA12878 | ONT Canu assembly | ftp://ftp.ncbi.nlm.nih.gov/genomes/all/GCA/900/232/925/GCA_900232925.2_NA127878-rel5/GCA_900232925.2_NA127878-rel5_genomic.fna.gz |
| *H. sapiens* NA12878 | Reference Genome | ftp://ftp.ncbi.nlm.nih.gov/genomes/all/GCF/000/001/405/GCF_000001405.39_GRCh38.p13/GCF_000001405.39_GRCh38.p13_genomic.fna.gz |

Table S3. Summary of stLFR and NGS datasets used in this work

| | Number of Read pairs | Average insert size(bp) | Length of read pair (bp) | Coverage | Number of Barcode |
|---|---|---|---|---|---|
| stLFR dataset of NA12878 chr19 | 21,507,205 | 170 | 100 | 73.4 | 1,105,848 |
| stLFR dataset of NA12878 whole genome sequences (WGS) | 1,035,984,388 | 229 | 100 | 64.6 | 37,196,043 |
| NGS data sets of NA12878 WGS | 479,007,051 | 390 | 150 | 47.9 | 0 |

Table S4. Control parameters used in different scaffolders for different datasets.

| | Chr19 MaSuRCA Contig | Human NA12878 MaSuRCA Contig | Human NA12878 SOAPdenovo2 Scaffold | Human NA12878 ONT Contig |
|---|---|---|---|---|
| SLR-superscaffolder | -MB 7000, -HB 3500, -T 0.1, -P 200 | -MB 7000, -HB 3500, -T 0.1, -P 1000 | -MB 7000, -HB 3500, -T 0.1, -P 300 | -MB 7000, -HB 3500, -T 0.1, -P 200 |
| fragScaff | -E 20000, -j 5, -u 2, -p=A | -E 20000, -j 5, -u 2, -p=A | -E 20000, -j 5, -u 2, -p=A | -E 20000, -j 5, -u 2, -p=A |
| Architect | -T 3, -E 0.15, -P 0.1 | -T 3, -E 0.15, -P 0.1 | -T 3, -E 0.15, -P 0.1 | -T 3, -E 0.15, -P 0.1 |
| ARKS | -k 100, -a 1.0 | -k 30, -a 1.0 | -k 30, -a 1.0 | -k 30, -a 1.0 |

Table S5. Evaluation of Chr19 assemblies based on MaSuRCA contigs by different scaffolders with the optimal parameters after the parameter sweeps. Scaffolding of the draft assembly was performed with SLR-superscaffolder (MST_BIN_SIZE, HT_BIN_SIZE, CLUSTER and PE_SEED_MIN abbreviated to "MB", "HB", "T" and "P" respectively), fragScaff (-m 1 -C 10 -t 8), Architect (--rc-abs-thr, --rc-rel-edge-thr and --rc-rel-prun-thr abbreviated to "T", "E" and "P" respectively), and ARKS -t 8 -c 5 -j 0.5 -z 3000 -e 30000 -m 50-6000 -r 0.05).

*Note*: The runtime for MaSuRCA is the total time for whole assembling process from raw reads to final scaffolds.

| | Control parameters | Number of scaffolds(>1000bp) | Largest scaffold (bp) | Total assembled length (bp) | NG50 (bp) | NGA50 (bp) | Relocation | Inversion | Number of misassemblies | Time | Peak Memory |
|---|---|---|---|---|---|---|---|---|---|---|---|
| SLR-superscaffolder | -MB 7000, -HB 3500, -T 0.1, -P 200 | 1,461 | 13,595,631 | 62,941,506 | 8,696,596 | 873,719 | 160 | 9 | 169 | 56min | 1.51GB |
| fragScaff | -E 20000, -j 5, -u 2, -p=A | 1,611 | 1,789,170 | 73,536,076 | 343,719 | 25,384 | 2,264 | 4 | 2,268 | 4h54min | 1.42GB |
| Architect | -T 3, -E 0.15, -P 0.1 | 3,286 | 179,888 | 57,015,198 | 29,949 | 25,783 | 465 | 4 | 469 | 4h49min | 1.42GB |
| ARKS | -k 100, -a 1.0 | 2,596 | 2,452,186 | 56,780,331 | 196,104 | 33,812 | 749 | 28 | 777 | 17min | 4.98GB |
| MaSuRCA | GRAPH_KMER_SIZE = 63, cgwErrorRate=0.15 | 3,499 | 148,918 | 57,069,141 | 27,512 | 26,350 | 157 | 5 | 162 | 3h28min | 17.02GB |

Table S6. Evaluation of Chr19 assemblies for different tests.

| | | Number of scaffolds (>1000 bp) | Largest scaffold (bp) | Total assembled length (bp) | NG50 (bp) | NGA50 (bp) | Relocation | Inversion | Number of misassemblies |
|---|---|---|---|---|---|---|---|---|---|
| MaSuRCA | | 3,779 | 148,918 | 57,069,141 | 27,512 | 26,350 | 157 | 5 | 162 |
| SLR-superscaffolder | | 2,186 | 13,595,631 | 62,941,506 | 8,696,596 | 873,719 | 160 | 9 | 169 |
| | No locally scaffolding | 2,213 | 13,621,100 | 63,116,164 | 8,706,614 | 850,387 | 168 | 15 | 183 |
| SLR-superscaffolder (with different length thresholds bp) | 1,000 | 1,329 | 23,756,072 | 76,490,713 | 11,754,257 | 110,175 | 990 | 207 | 1197 |
| | 3,000 | 1,491 | 22,236,846 | 65,689,522 | 10,200,611 | 276,992 | 479 | 40 | 519 |
| | 5,000 | 1,900 | 22,599,625 | 62,039,221 | 10,076,573 | 847,484 | 189 | 27 | 216 |
| | 7,000 | 2,186 | 13,595,631 | 62,941,506 | 8,696,596 | 873,719 | 160 | 9 | 169 |
| | 10,000 | 1,920 | 11,703,047 | 64,839,557 | 4,047,100 | 740,961 | 144 | 6 | 150 |
| | 15,000 | 3,218 | 2,970,432 | 65,941,210 | 803,638 | 286,442 | 137 | 4 | 141 |
| | 20,000 | 3,634 | 1,944,702 | 63,349,858 | 241,262 | 88,273 | 125 | 6 | 131 |

Table S7. Statistics of tip and long junctions before and after conducting the screening algorithm.

| | Tip junction | | Long junction | |
|---|---|---|---|---|
| For the node | Unique | Non-unique | unique | Non-unique |
| Before | 1375 | 451 | 105 | 249 |
| After | 1049 | 100 | 0 | 0 |

Table S8. Statistics of local properties of tip and long junctions before and after conducting the screening algorithm.

| | Tip junction | | Long junction | |
|---|---|---|---|---|
| For local graph of the node | Unique | Non-unique | unique | Non-unique |
| Before | 1045 | 781 | 41 | 313 |
| After | 917 | 232 | 0 | 0 |

Table S9. Runtime statistics for SLR-superscaffolder step by step.

| | MaSuRCA contigs | | SOAP*denovo* scaffolds | | ONT contigs | |
|---|---|---|---|---|---|---|
| | Wall-clock time(h) | Percentage (%) | Wall-clock time (h) | Percentage (%) | Wall-clock time(h) | Percentage (%) |
| Total | 56.1 | 100.0 | 42.1 | 100.0 | 104.6 | 100.0 |
| Data preparing | 31.2 | 55.6 | 33.4 | 79.4 | 41.8 | 39.9 |
| Ordering | 12.0 | 21.4 | 2.5 | 6.0 | 1.8 | 1.7 |
| Orientating | 7.8 | 13.9 | 1.1 | 2.7 | 1.2 | 1.2 |
| Local scaffolding | 1.7 | 3.1 | 2.7 | 6.4 | 1.4 | 1.3 |
| Estimating gap size | 3.4 | 6.1 | 2.3 | 5.6 | 58.5 | 55.9 |

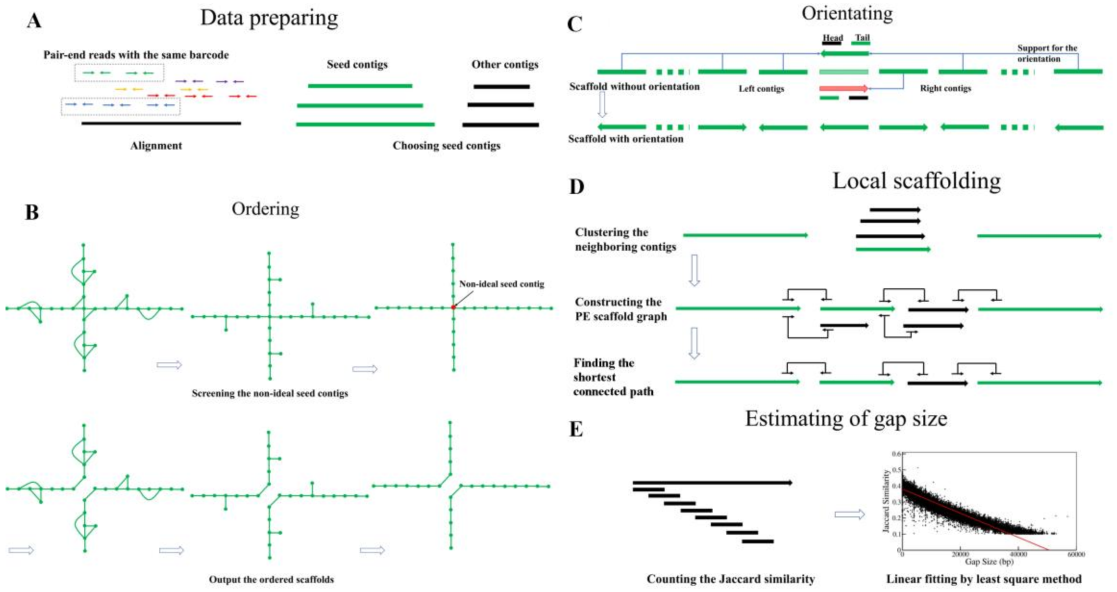


Figure S1. Overall scheme of SLR-superscaffolder. (A) In data preparing, two sub-processes are included: aligning stLFR reads to the draft assembly and choosing seed contigs. (B) In ordering, suspicious seed contigs are interactively screened as shown in the upper three figures, and then ordered scaffolds are generated as shown in the lower three figures. (C) In orientating, $n_{th}$-order neighboring contigs in each ordered scaffold are used to determine the orientation state of each contig. (D) In local scaffolding, contigs near a paired neighboring contigs in the scaffold are firstly clustered by co-barcoding information, and then a scaffold graph is further constructed by PE information. Finally, the shortest path among the neighboring contigs are determined as local scaffolds. (E) In the estimating of gap size, a statistical relation between Jaccard similarity of shared barcodes and distance is calculated for long contigs, and then an approximately linear relation is fitted using the least square method.

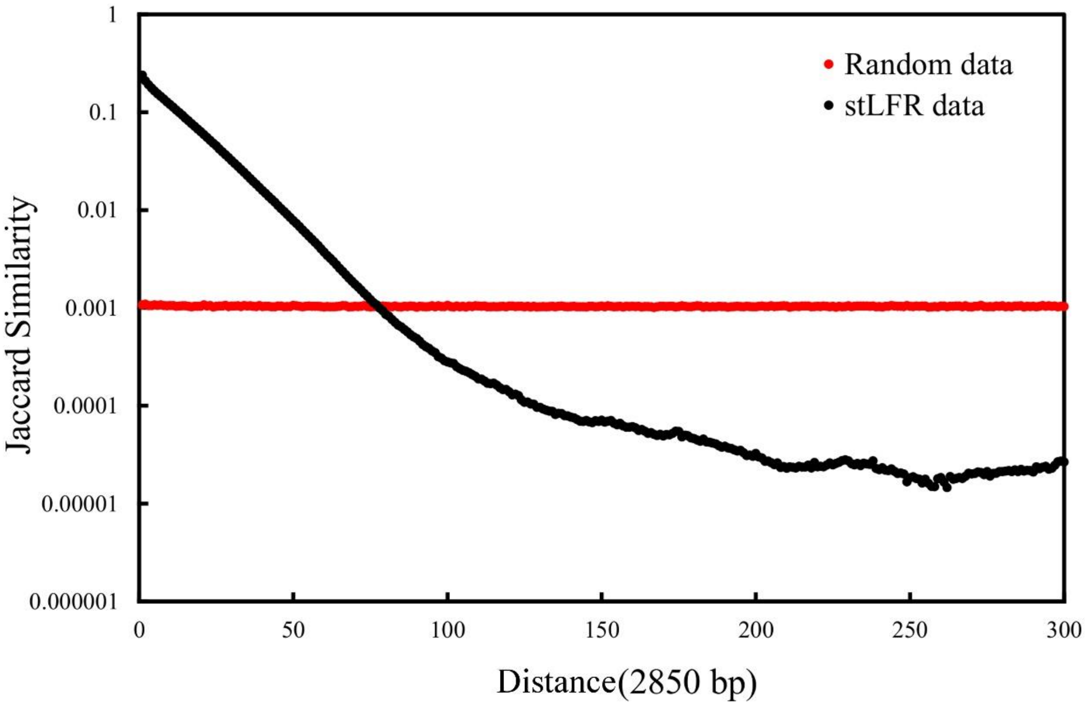


Figure S2. Relations between Jaccard similarity of barcodes and distance for two sequences in the reference for stLFR reads and randomly barcoded reads. The randomly barcoded reads are generated by randomly distributing the stLFR barcodes to the same read sets.

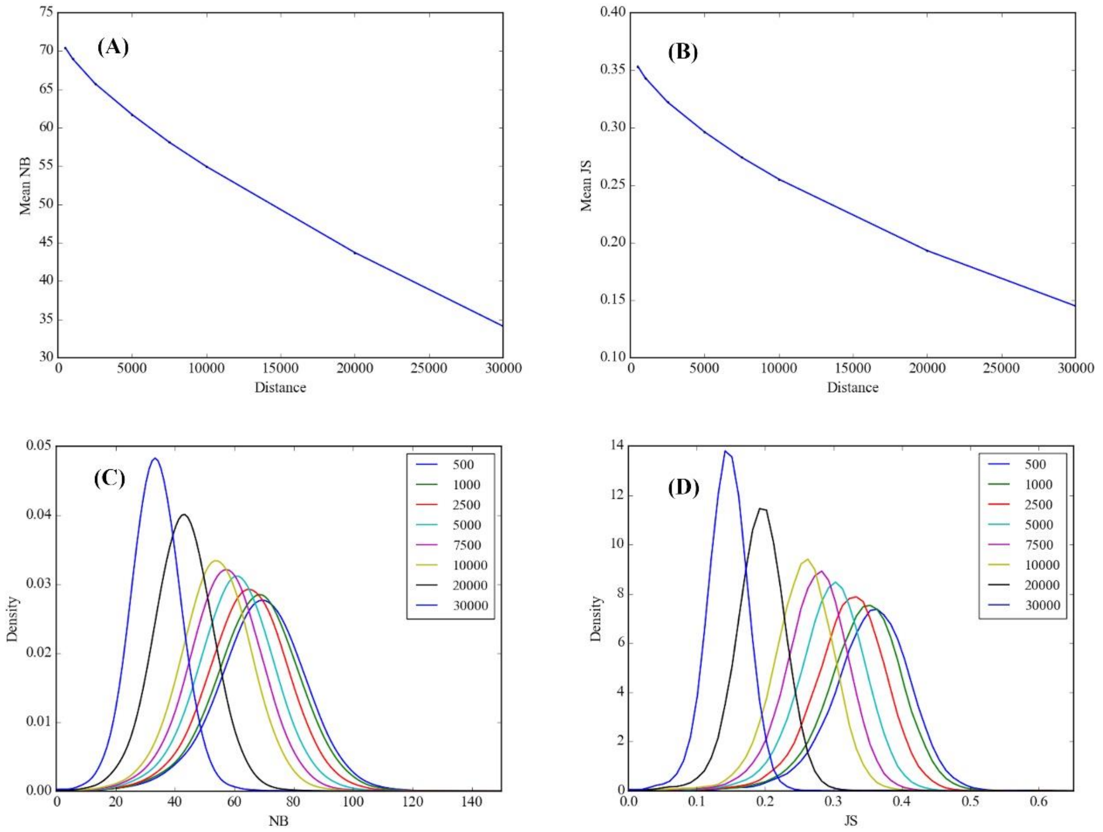


Figure S3. Mean NB of two bins at different distances (A), mean JS of two bins at different distances (B), distribution of NB of two bins at different distances (C), distribution of JS of two bins at different distances (D) with bin size of 1200 bp.

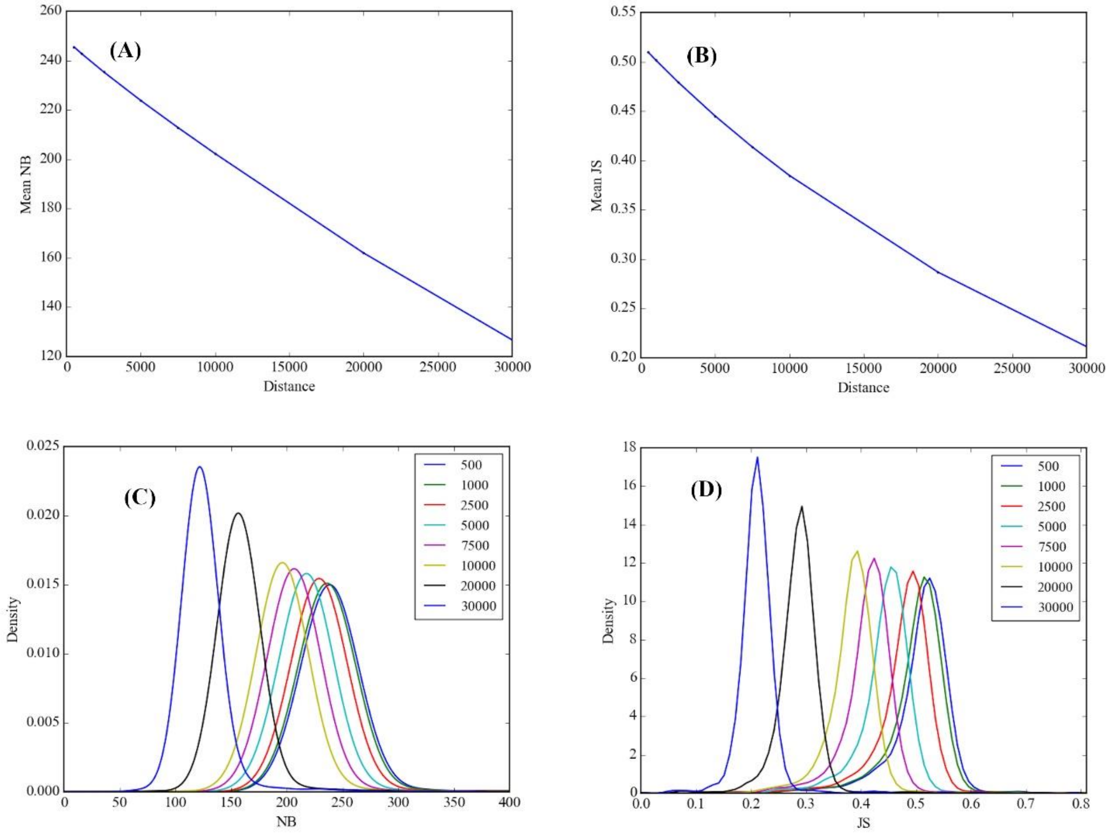


Figure S4. Mean NB of two bins at different distances (A), mean JS of two bins at different distances (B), distribution of NB of two bins at different distances (C), distribution of JS of two bins at different distances (D) with bin size of 20000 bp.
